# Sex-Dependent Responses in Mice to Indomethacin-Induced Injury and Gut Microbiome-Targeted Alleviation

**DOI:** 10.1101/2025.04.25.650491

**Authors:** Jianan Zhang, Josh J. Sekela, Jun Yang, Rani S. Sellers, Aadra P. Bhatt, Matthew R. Redinbo

## Abstract

Nonsteroidal anti-inflammatory drugs (NSAIDs) are used widely but produce gastrointestinal (GI) toxicities in both short- and long-term users. Previous studies have shown that the intestinal microbiota play an important role in gut damage and that gut microbial β-glucuronidase (GUS) inhibitors can alleviate NSAID-induced injury in male mice by blocking the GI reactivation of NSAID-glucuronides. Here, in both male and female C57BL/6 mice, we examine the effects of indomethacin alone and with the GUS inhibitor UNC10201652. Oral delivery of 5 mg/kg body weight indomethacin over five days decreased body weight, induced colonic and hepatic inflammatory cytokine gene expression, and enlarged the spleens of both male and female mice. However, sex-specific inflammatory responses to indomethacin were observed, with males demonstrating more colonic injury while females presented greater splenic and hepatic toxic responses. Females also showed a unique indomethacin-induced bloom of fecal Verrucomicrobia as measured by *16S* rRNA metagenomic sequencing. UNC10201652 alleviated aspects of these indomethacin-induced toxicities, including features of the male-specific colonic damage and the female-specific compositional changes and spleen and liver toxicities. Thus, GI and non-GI tissues in male and female mice respond distinctly to indomethacin-induced damage. These findings advance our understanding of how sex impacts systemic responses to xenobiotic exposure and may lead to improved therapeutic outcomes with these widely used drugs.

## INTRODUCTION

Non-steroidal anti-inflammatory drugs (NSAIDs) are critical to health but present significant problems. NSAIDs treat pain, fever and inflammation are among the most widely used drugs in the world, both over-the-counter and by prescription ^1,2^. However, NSAIDs cause lower gastrointestinal (GI) toxicities in short- and long-term users. The mechanisms of NSAID-induced gut toxicity are multifactorial, and arise from both topical effects on the intestinal epithelial layer and basolateral effects subsequently produced by host immune responses ^3^. The topical effects include the decoupling of mitochondrial function in epithelial cells, which decrease ATP production and generate reactive oxygen species ^3^. They also include electrophilic stress and endoplasmic reticulum stress responses that together enhance cell death and increase intestinal permeability ^3^. Finally, the inhibition of the COX-dependent generation of prostaglandin E2 (PGE2) further decreases the restitution of the epithelial layer and promotes the erosion of this barrier along the GI tract ^4^. Indeed, NSAID-induced lower GI bleeds and ulcerations are estimated to cause 16,500 deaths in the United States each year ^5,6^.

It has been shown that the gut microbiota play an important role in the intestinal toxicities of NSAIDs. Germ-free rodents and rodents receiving antibiotics do not develop lower GI ulcers when treated with the NSAID ^7^. Furthermore, a specific set of gut microbial enzymes has been linked to exacerbating NSAID-induced lower intestinal ulcers, loss of epithelial barrier function, and poor outcomes after intestinal surgery ^8,9^. NSAIDs reach the GI tract as inactive glucuronide conjugates produced by host protective processes. However, gut microbial β-glucuronidase (GUS) enzymes have been shown to efficiently remove the glucuronide sugar to regenerate the active and anionic forms of the drugs in the lumen of the GI tract ^10^. These effects have been blocked in rodent models by the use of gut microbial GUS inhibitors that remain in the GI tract and are selective for bacterial GUS proteins without inhibiting the host GUS, which is expected to be essential ^10^. Use of GUS inhibitor has blocked lower intestinal ulcers in mice treated with diclofenac, indomethacin and ketoprofen ^3,10–12^. GUS inhibition has also been shown to prevent anastomosis caused by diclofenac in a rat surgery model ^13^. Taken together, these results demonstrate that the gut microbiota facilitate poor lower intestinal outcomes with NSAIDs by producing reactivated drug in the GI lumen that drive the topical effects leading to ulcers and loss of barrier function.

While gut microbial GUS enzymes have been shown to reactivate a range of small molecule therapeutics in the mammalian GI tract, they also play critical roles in processing hormones, neurotransmitters and other endobiotics that reach the gut as glucuronide conjugates. These processes are part of enterohepatic recirculation important for homeostasis, but have been hypothesized to contribute to hormone-driven diseases like breast cancer ^14^. Specific human microbial GUS proteins have been shown to efficiently convert the inactive glucuronide conjugates of estradiol and estrone back to active hormones in both in vitro assays and using proteins identified from multi-omics studies ^15,16^. Therefore, it is important to consider the gut microbiota’s role in NSAID-induced toxicity in the context of sex as a biological variable. In addition, sex is important in the human use of NSAIDs because nearly 80% of arthritis patients employ these therapeutics and the female to male ratio for this disease is 3-to-1 ^17,18^. In other words, women consume NSAIDs more frequently than men. Indeed, a study of 4.6 million people showed that women fill more NSAID prescriptions than men and that overall NSAID use increases with age ^19^.

NSAIDs produce sex-dependent differences in humans. Healthy women typically show higher fecal microbiota diversity than healthy men, indomethacin was found to decrease this diversity in women more than men ^20^. Estrogens also appear to protect the gut epithelial layer and may lessen NSAID-induced topical damage in females compared to males ^21^. Finally, differences in mucin secretion and prostaglandin synthesis between men and women may also contribute to differential NSAID-induced gastrointestinal damage ^22^. Because sex-dependent differences in NSAID responses have been observed in humans, we chose to examine the effects of sex and gut microbial GUS inhibition on the development of indomethacin-induced toxicities in both male and female mice. To date, published work on NSAID-induced damage and gut microbial GUS inhibition has examined only male rodents^3,10–12^. Additionally, those studies employed a first-generation gut microbial GUS inhibitor and not the second-generation molecule UNC10201652 that has been shown blocked intestinal damage caused by the anti-cancer drug irinotecan ^23^ and the consumer products antimicrobial agent triclosan ^24^. Here, we examine indomethacin-induced damage in both male and female C57BL/6 mice with and without UNC10201652. We find sex-specific differences in response to indomethacin, with males exhibiting more intestinal damage and females demonstrating increased hepatic and splenic toxicities. We also find that indomethacin induces a bloom of Verrucomicrobia in the fecal microbiota of female mice. Finally, we show that UNC10201652 alleviates measures of male- and female-specific toxicities, along with the gut microbial compositional changes in female mice. Taken together, these results reveal specific sex-dependent differences in response to indomethacin in mice, results that and may eventually improve NSAID use in humans.

## RESULTS

### Indomethacin in Male and Female Mice

We randomly separated acclimatized male and female C57BL/6 into two groups of 5-6 animals per group and delivered 5 mg/kg body weight indomethacin or vehicle by oral gavage daily for five days (**Figure 1A**). Mice were weighed daily. Animals receiving indomethacin exhibited significantly reduced weight compared to those receiving vehicle (**Figure 1B**). After five days of indomethacin or vehicle, mice were sacrificed and examined for evidence of drug-induced damage. In the colon, mice receiving indomethacin exhibited epithelial damage and ulcers (**Figure 1C**). Specifically, two of five female indomethacin mice developed ulcers, while five of the six male mice showed ulcers with indomethacin, a level of damage that reached statistical significance (**Figure 1C**). By these first two metrics, male and female mice responded in a similar manner to indomethacin. In terms of immune cells, though, only male mice exhibited increased colonic infiltrates with indomethacin, while female mice failed to show this effect (**Figure 1D**). Male and female mice both exhibited significantly increased *Il-1β* and *Mcp-1* inflammatory cytokine expression with indomethacin in the colon compared to vehicle controls (**Figure 1E**). However, with indomethacin, female mice showed increased colonic *Muc2* expression, while male mice demonstrated decreased expression of *Tff3* (**Figure 1F**). Thus, five days of oral 5 mg/kg indomethacin reduces body weight, and leads to colonic damage, ulcers and similar levels of inflammatory cytokine expression in both male and female mice, while differences were seen in colonic immune cell infiltrates as well as *Muc2* and *Tff3* expression.

**Figure 1.**
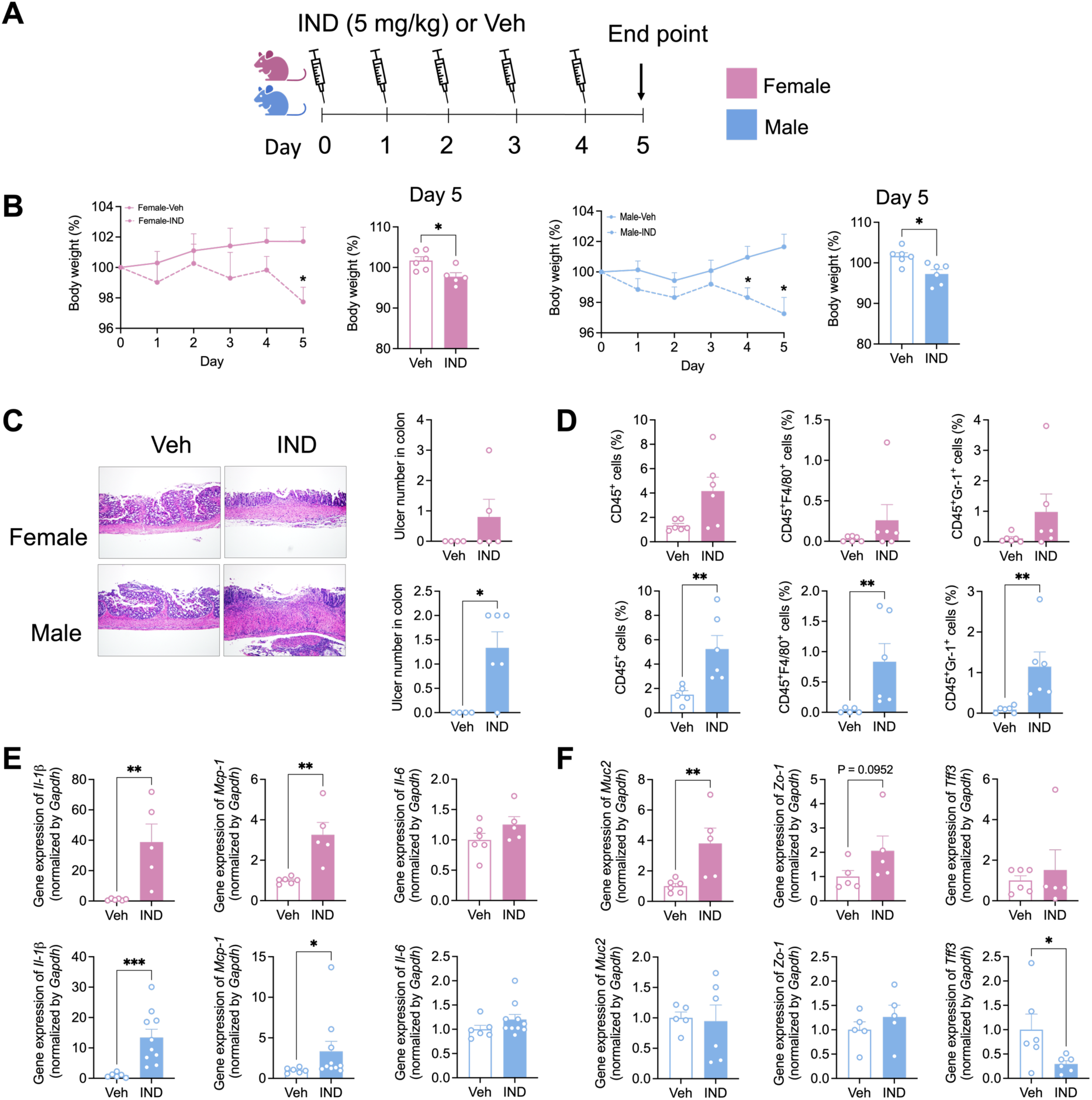
The effect of indomethacin in colon in both sexes of mice. **A.** C57BL/6 mice were treated with 5 mg/kg dose of indomethacin or veh via oral gavage for both male and female mice from day 0 to day 4, and the end point is at day 5. B. Percentage of body weight. C. Indomethacin induced ulceration in colon. Left: representative H&E histological images of colon (200X). Right: ulcer numbers in the colon. D. Immune cell infiltration in colon: CD45^+^ cells, CD45^+^F4/80^+^ cells and CD45^+^Gr-1^+^ cells. E-F. Gene expression of pro-inflammatory cytokines and intestinal barrier integrity markers in the colon. The data are mean ± SEM. *P < 0.05, **P < 0.01, ***P < 0.001.

We next examined the liver and spleens of these animals. We found that indomethacin induced splenic enlargement in both sexes of mice (**Figure 2A**). However, in the liver, only male mice receiving indomethacin exhibited bacterial infiltration, which also extended to the blood of male mice (**Figures 2B-D**). By contrast, female mice did not show significant increases in liver or blood bacterial levels (**Figures 2B-D**). Both male and female mice receiving indomethacin showed increased liver *Il-1β* cytokine expression (**Figure 2E**), as well as expression of *Cd36* factor associated with liver damage (**Figure 2F**). Liver expression of *Ppar-α*, a regulator of hepatic lipid oxidation, was found to be decreased only in female mice (**Figure 2F**). Thus, indomethacin enlarges the spleen and damages the liver of both male and female mice, but only male mice showed evidence of bacterial infiltration in the liver and blood.

**Figure 2.**
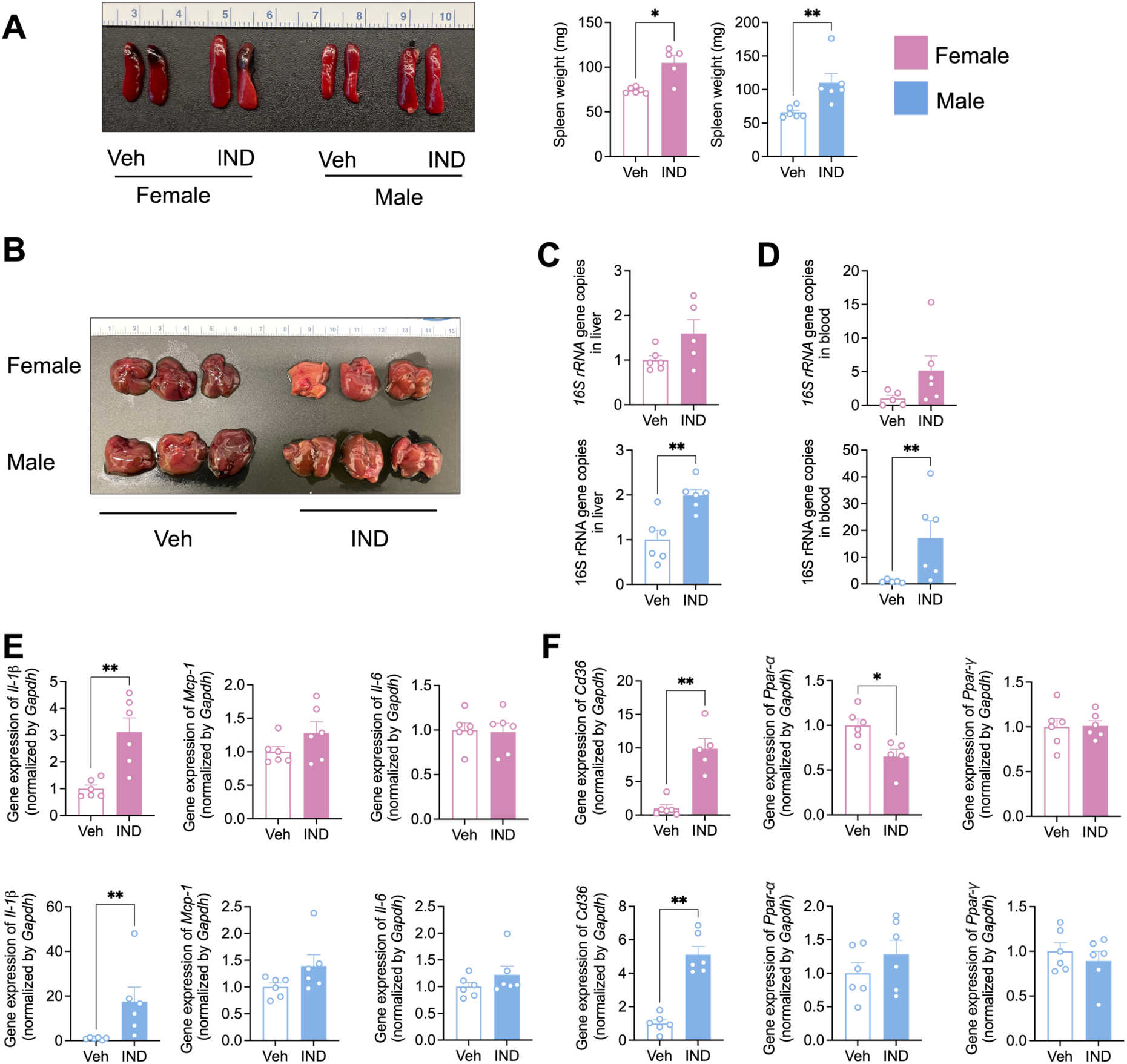
The effect of indomethacin in liver and spleen in both sexes of mice. **A.** Spleen weight. **B.** Indomethacin induced injury in liver. **C.** Bacteria translocation in liver. **D.** Bacteria translocation in blood. **E.** Gene expression of pro-inflammatory cytokines in liver. **F.** Gene expression of fatty liver markers in liver. The data are mean ± SEM. *P < 0.05, **P < 0.01, ***P < 0.001.

Finally, in the ileum, we found little evidence for epithelial damage and ulcers with indomethacin in either sex (**Figure S1A**). However, female mice that received indomethacin showed significantly higher of CD45^+^ cell infiltrates, while male mice showed significantly higher CD45^+^Gr-1^+^ cell (**Figure S1B**). Like the colon and liver, though, both male and female mice showed significantly increased *Il-1β* compared with indomethacin (**Figure S1C**). Thus, 5 mg/kg oral indomethacin for five days produces several common features of damage to the colon, liver and ileum in both male and female mice. However, sex-dependent differences were observed with indomethacin in immune cell infiltrates, and only male mice exhibited bacteria in the liver and blood post treatment.

### Indomethacin and UNC10201652 (GUSi) in the Male and Female Mouse Colon

We next examined the effects of UNC10201652 (GUSi) in alleviating indomethacin induced damage in mice. Three groups of male and female mice were established. One group of each sex received 2 mg/kg GUSi by oral gavage for three days, while the other two groups received a control treatment. One control group continued receiving the control treatment, followed by a vehicle for an additional five days. The second control group also received the control treatment but was given 5 mg/kg indomethacin by oral gavage for five days. The GUSi-treated group continued receiving GUSi and was additionally given 5 mg/kg indomethacin for five days (**Figure 3A**). Control or GUSi were provided in the morning, while indomethacin or vehicle were provided in the evening.

**Figure 3.**
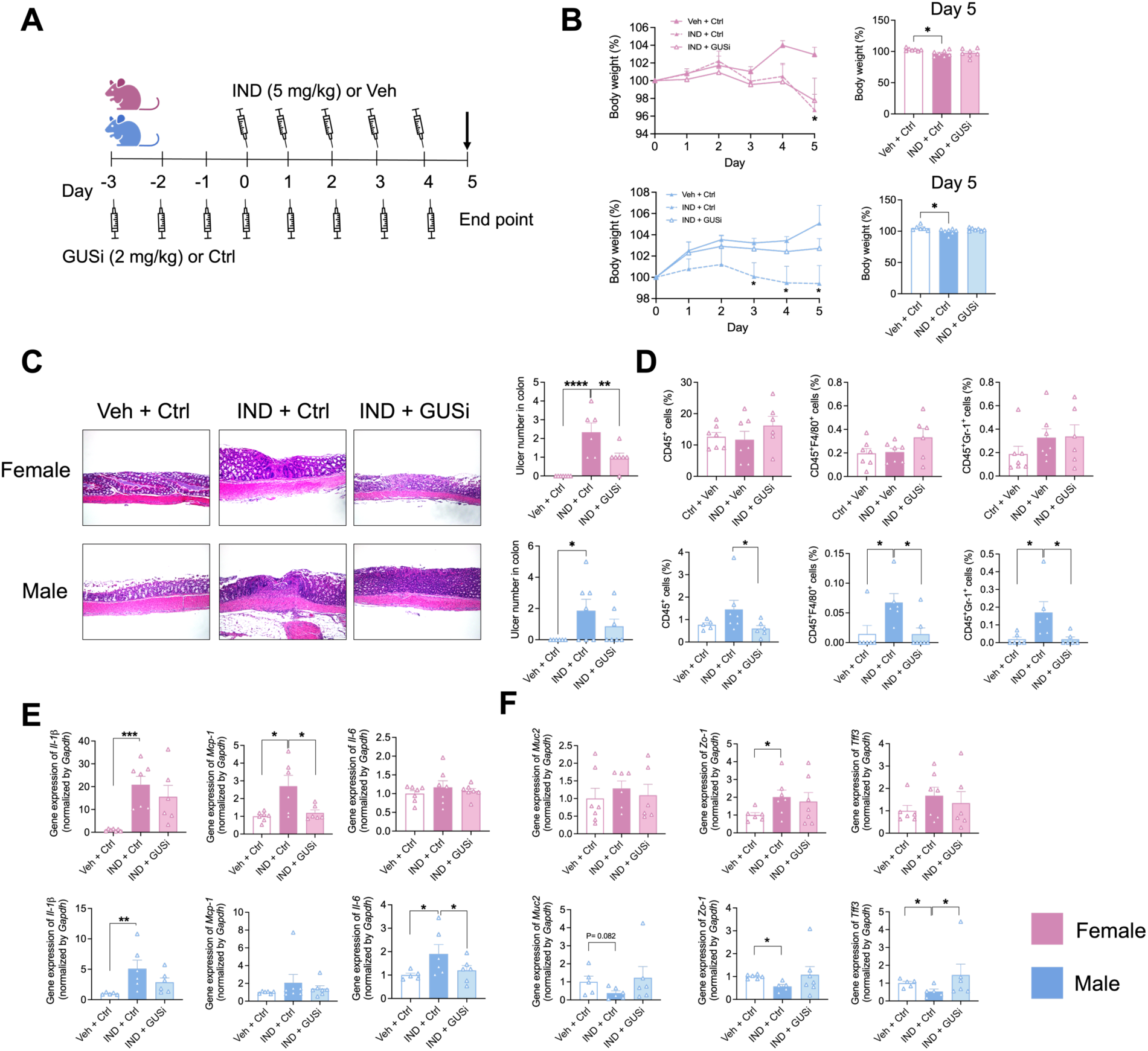
The effect of inhibition microbial β-glucuronidase (β-GUS) enzymes in indomethacin-induced colon damages in both sexes of mice. **A.** C57BL/6 mice were treated with 2 mg/kg b.w. of microbial β-glucuronidase (β-GUS) enzymes inhibitor (GUSi, UNC10201652) or ctrl via oral gavage from day -3 to day 4; and stimulated with 5mg/kg b.w. of indomethacin or veh per day from day 0 to day 4. The end point of this study is at day 5. **B.** Percentage of body weight. **C.** Indomethacin induced ulceration in colon. Up: representative H&E histological images of colon (200X). Down: ulcer numbers in colon. **D.** Immune cell infiltration in colon: CD45^+^ cells, CD45^+^F4/80^+^ cells and CD45^+^Gr-1^+^ cells. **E-F.** Gene expression of pro-inflammatory cytokines and intestinal barrier integrity markers in the colon. The data are mean ± SEM. *P < 0.05, **P < 0.01, ***P < 0.001.

Like the data shown in **Figure 1**, male and female mice receiving indomethacin showed significantly reduced body weight after 3-5 days (**Figure 3B**). GUSi had no effect on body weight of female mice but did show a trend in improving the body weight of the male mice (**Figure 3B**). All mice receiving indomethacin in the absence of GUSi showed colonic epithelial damage and ulcers (**Figure 3C**). Female mice receiving GUSi showed significantly reduced ulcers compared to indomethacin plus control, indicating that GUSi has the potential to reduce colonic ulceration in female mice (**Figure 3C**). Male mice, by contrast, did not show a difference in colonic ulcers with GUSi (**Figure 3C**).

In terms of immune cell infiltrates in indomethacin-treated mice, only male mice showed significant increases in colonic CD45^+^F4/80^+^ cells and CD45^+^Gr-1^+^ cells, while female mice showed no evidence of immune cell infiltrates in the colon (**Figure 3D**). However, while GUSi did not block colonic ulcers in males, it did reduce colonic immune cell infiltrates (CD45^+^ cells, CD45^+^F4/80^+^ cells and CD45^+^Gr-1^+^ cells) in indomethacin-treated animals (**Figure 3D**). Thus, male mice respond to indomethacin by increasing immune cell infiltrates in the colon, a response distinct from females, and GUSi was found to block these infiltrates in male mice.

Mice of both sexes showed increased colonic *Il-1β* expression with indomethacin and no reduction of this expression when GUSi was also administered (**Figure 3E**). However, only female mice showed increased colonic *Mcp-1* expression with indomethacin, an effect that was blunted by GUSi (**Figure 3E**). By contrast, male mice showed increased colonic *Il-6* expression with indomethacin, an increase that was suppressed by GUSi (**Figure 3E**).

We next examined the colonic expression of *Muc2*, *Zo-1* and *Tff3*. We found that female mice showed an increase in *Zo-1* with indomethacin, indicative of improved colonic epithelial tight junctions in these animals (**Figure 3F**). By contrast, we found that male mice showed a decrease in expression of *Zo-1, Muc2* and *Tff3* with indomethacin, all indicative of decreased colonic epithelial integrity (**Figure 3F**). These results may provide an explanation for the increased liver and blood bacteria noted above in male mice post indomethacin (**Figures 2B-D**). GUSi normalized *Tff3* expression in male mice and led to trends in increased *Muc2* and *Zo-1* expression, as well (**Figure 3F**). Thus, indomethacin generates sex-dependent differences in colonic damage in mice, and GUSi similarly shows evidence sex-dependent differences in its ability to block these effects.

### Indomethacin and UNC10201652 (GUSi) in Non-Colonic Regions of Male and Female Mice

In the ileum, we observed a trend toward increased epithelial damage and ulcers with indomethacin (**Figure S2A**). Unlike the colon, however, neither male nor female mice showed increased immune cell infiltration with indomethacin, although GUSi decreased CD45^+^F4/80^+^ levels in female mouse ileum (**Figure S2B**). Inflammatory cytokine expression was not changed in male ileum with indomethacin. Female mouse ileal regions, however, showed increases in both *Il-1β* and *Mcp-1* with indomethacin (**Figure S2C**). Similarly, indomethacin led to changes in epithelial integrity in the female ileum, as expression of *Zo-1* and *Tff3* significantly increased, with Tff3 levels effectively reduced with the addition of GUSi (**Figure S2D**). By contrast, these factors were not changed in the male ileum with indomethacin alone or with GUSi (**Figure S2D**). Thus, 5 mg/kg oral indomethacin for five days produces moderate sex-dependent differences in the mouse ileum.

While male and female mice both exhibited increased blood bacteria, the magnitude of the effect was 2-fold higher in male mice and bacterial infiltration into the blood was blocked by GUSi only in male animals (**Figure 4A**). The spleens of both male and female mice increased in weight with indomethacin, but GUSi was able to block this increase only in male mice (**Figure 4B**). We next examined splenic inflammatory cytokine expression. We found that only female mice showed increased *Il-1β* and *Il-6* expression with indomethacin, and that GUSi blocked *Il-1β* expression but increased *Il-6* expression in these animals (**Figure 4C**). By contrast, male mice showed no change in splenic *Il-1β, Mcp-1* and *Il-6* expression with indomethacin (**Figure 4C**). Thus, significant differences in male and female mice are observed in splenic cytokine responses with indomethacin and GUSi.

**Figure 4.**
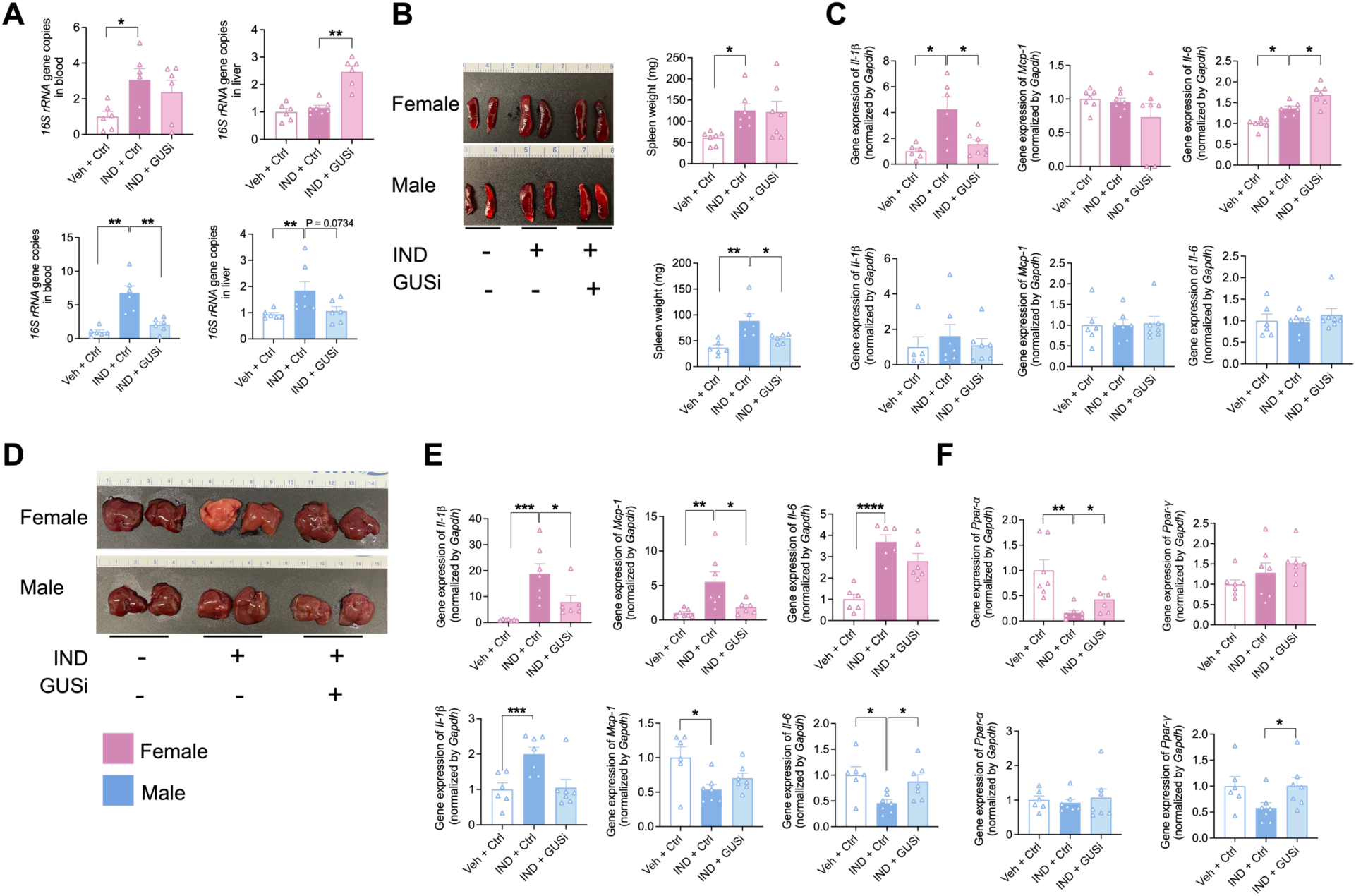
The effect of inhibition microbial β-glucuronidase (β-GUS) enzymes in indomethacin-induced organ injury in both sexes of mice. **A.** Bacteria translocation in blood. **B.** Spleen weight. **C.** Gene expression of pro-inflammatory cytokines in spleen. **D.** Indomethacin induced injury in liver. **E.** Gene expression of pro-inflammatory cytokines in liver. **F.** Gene expression of lipid accumulation in liver. The data are mean ± SEM. *P < 0.05, **P < 0.01, ***P < 0.001.

In this study, female livers again exhibit more damaged with indomethacin than males, and the female livers were protected by GUSi (**Figure 4D**). Furthermore, while liver *Il-1β* expression was significantly increased with indomethacin in both male and female mice, the magnitude of the effect was ∼10-fold higher in female animals and was significantly blocked by GUSi (**Figure 4E**). We also observed sex-dependent differences in liver *Mcp-1* and *Il-6* expression with both indomethacin and GUSi. Female mice showed increased *Mcp-1* and *Il-6* expression, while male mice showed expression of both factors, and GUSi reduced *Mcp-1* expression in the female liver while increasing *Il-6* expression in the males (**Figure 4E**). Finally, we examined the expression of markers of fatty liver disease. Female mice showed a decrease in liver *Ppar-α* expression with indomethacin, and an improvement with GUSi (**Figure 4F**). By contrast, male mice showed a trend toward decreased *Ppar-γ* expression with indomethacin, and GUSi significantly increased the expression of this factor over levels seem with indomethacin alone (**Figure 4F**). Thus, sex-dependent differences were observed in the effects of indomethacin and GUSi on markers of inflammation and fat accumulation in the mouse liver.

### Indomethacin and Indomethacin-Glucuronide Levels in Male and Female Mice

We next examined the levels of indomethacin and indomethacin-glucuronide in mice after five days of oral 5 mg/kg indomethacin with and without GUSi. Previous data have shown that gut microbial β-glucuronidases can remove the inactivating glucuronic acid moiety and increase gut luminal levels active NSAID that cause toxicity (**Figure 5A**) but can be blocked by GUS inhibitors ^3,10–12^. We measured the levels of indomethacin and indomethacin-glucuronide in various compartments upon sacrifice. We found plasma levels of indomethacin showed no difference between male and female mice and were not affected by GUSi (**Figure 5B**). By contrast, plasma indomethacin-glucuronide levels were significantly higher (∼19 times) in the female animals and were not changed by GUSi (**Figure 5B**). We also observed that male and female mice showed no differences in the levels of indomethacin or indomethacin-glucuronide in the liver, indicating that the production of indomethacin-glucuronide was not altered in a sex dependent manner (**Figure 5C**). Male mice exhibited significantly higher indomethacin levels in the cecum and those levels were not altered by GUSi (**Figure 5D**), while no differences were found in the colons of either male or female animals (**Figure 5E**). Taken together, these results indicate that three days of pre-treatment with 2 mg/kg GUSi along with five further days of GUSi plus indomethacin did not impact active or inactive drug levels in this study. However, significant sex-dependent differences in plasma indomethacin-glucuronide and cecal indomethacin levels in female and male mice, respectively (**Figures 5B, D**), indicate that the routes of elimination of indomethacin-glucuronide formed in the liver differ in a sex-dependent manner. Female mice appear to traffic more indomethacin-glucuronide to the plasma for kidney elimination, while male mice appear to employ GI excretion via the bile duct.

**Figure 5.**
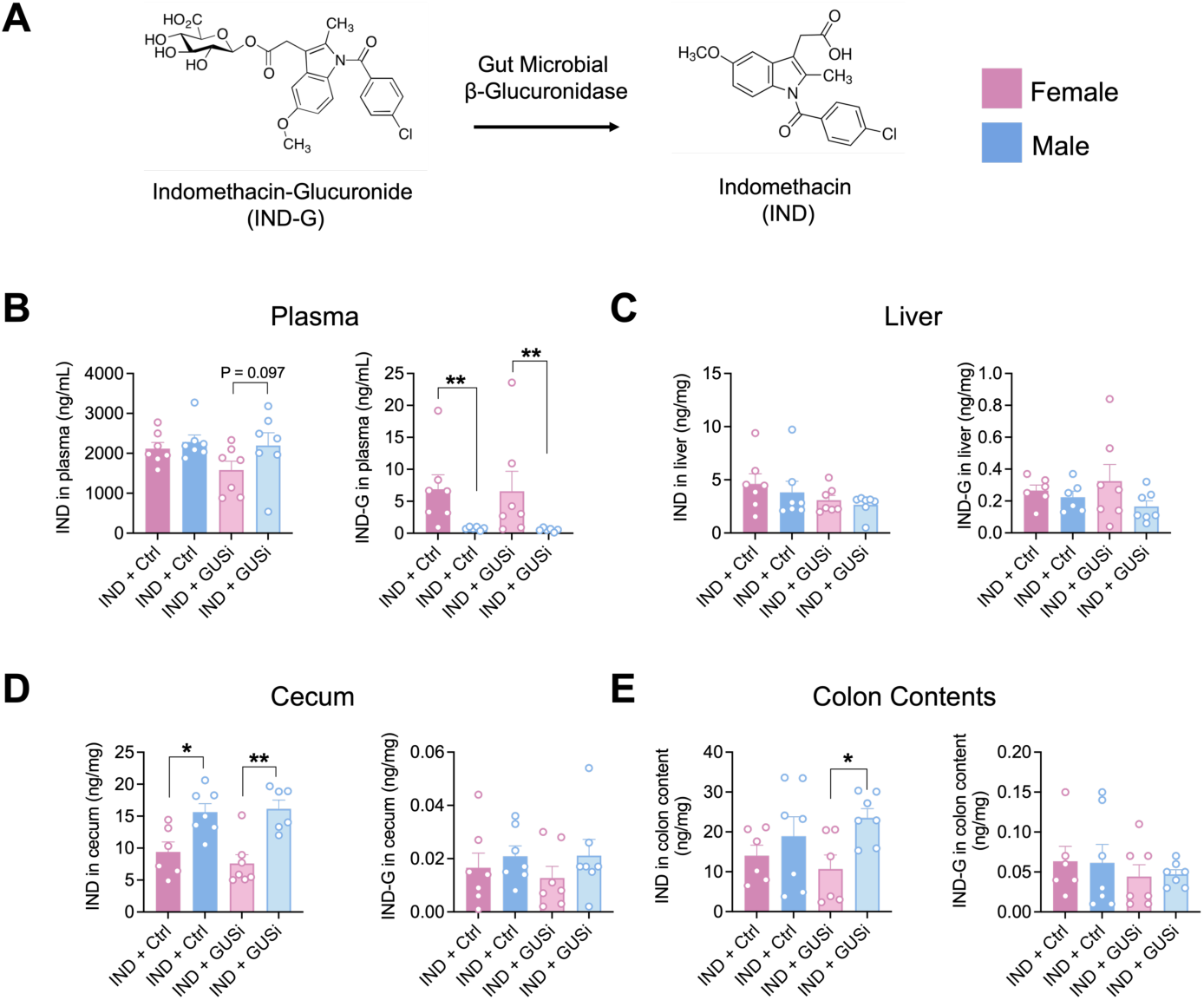
The indomethacin metabolites in plasma. **A.** Metabolites of indomethacin via microbial β-glucuronidase (GUS) enzymes. **B.** Level of indomethacin (IND) and indomethacin-Glucuronide (IND-G) in plasma. **C.** Level of IND and IND-G in liver. **D.** Level of IND and IND-G in cecum. **E.** Level of IND and IND-G in colon content. The data are mean ± SEM. *P < 0.05, **P < 0.01, ***P < 0.001.

### Gut Microbial Composition with Indomethacin in Male and Female Mice

Given the colonic damage produced by indomethacin in male and female mice, we examined the composition of the fecal microbiota using *16S rRNA* sequencing. We studied feces collected at day 0, after either GUSi or control but prior to indomethacin, and at day 5 after either indomethacin or vehicle (**Figure 6A**). We found that male mice exhibited higher gut microbial alpha diversity than female mice regardless of treatment with indomethacin or GUSi (**Figure 6B**). We further found that GUSi increased alpha diversity in female mice but only when used in conjunction with indomethacin (**Figure 6B**). Male and female mice were significantly different in terms of gut microbial beta diversity, and that sex drove the distinct clustering of the mice in our study by principal coordinate analysis (**Figure 6C**). The relative abundances of microbial phyla measured by *16S rRNA* sequencing at days 0 and 5 again showed that male and female mice start with distinct compositions, and that after indomethacin female mice show a unique increase in microbes from the Verrucomicrobia phylum (**Figure 6D**). Focusing on the most prevalent gut microbial phyla reveals that male mice have significantly more fecal Firmicutes taxa than females, while females exhibit higher levels of Bacteroidetes taxa than males (**Figure 6E**). Indomethacin and GUSi had no effects of those phyla, or on the Proteobacteria (**Figure 6E**). However, indomethacin treatment significantly increased the levels of Verrucomicrobia in the feces of female mice, but not male mice, and that this effect was significantly reduced when GUSi was present along with indomethacin (**Figure 6E**). We found that sex significantly alters microbial composition at the phylum, order, and genus levels. A complete comparison of the abundance of fecal microbiota by sex and by treatment at the phylum, order, and genus levels can be found in **Tables S2-S4**. Thus, the mice in our study started with distinct fecal microbiota compositions, and female mice uniquely showed a bloom of Verrucomicrobia in response to indomethacin treatment that can be reversed by GUSi. This observation provides additional evidence of sex-dependent differences in the effects of indomethacin in mice.

**Figure 6.**
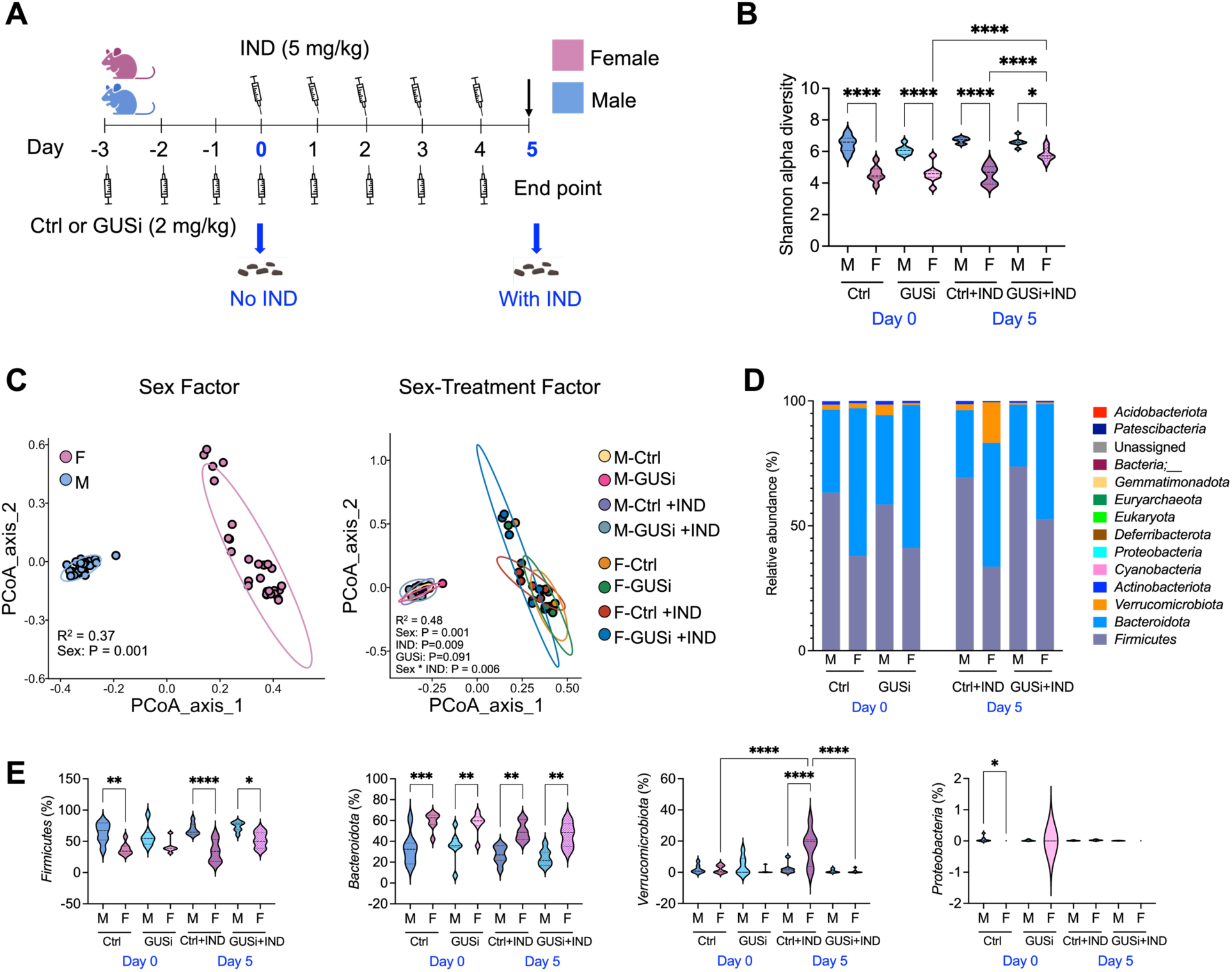
Inhibition of gut microbial GUS enzymes altered the gut microbiota in indomethacin-treated mice in both sexes. **A.** Fecal material was collected on day 0 (with no treatment of IND) and day 5 (with treatment of IND). **B.** Sex difference was showed in alpha diversity; male mice showed significantly higher diversity compared to female mice. **C.** Sex difference was showed in beta diversity; female mice showed diverse response to the treatment of IND and GUSi. **D.** IND and GUSi altered the composition of the microbiota at phylum level. **E.** Firmicutes, Bacteroidota, Verrucomicrobiota, and Proteobacteria showed significant difference among the groups. The data are mean ± SEM. *P < 0.05, **P < 0.01, ***P < 0.001, ****P < 0.0001.

## DISCUSSION

The oral delivery of 5 mg/kg body weight of the NSAID indomethacin for five days produced several toxicities in both male and female C57BL/6 mice, including weight loss, spleen enlargement, hepatic *Il-1β* expression, and colonic ulcers and inflammatory cytokine expression. However, sex-dependent differences were observed. Male mice showed colonic immune cell infiltration, loss of colonic epithelial integrity, and evidence of bacterial infiltration in both the liver and blood. Female mice showed higher splenic and liver inflammatory cytokine expression and an increase in colonic Verrucomicrobia with indomethacin. The levels of indomethacin-glucuronide were significantly higher in plasma of females while indomethacin levels were significantly higher in the cecum of males. This is the first report of sex-dependent differences in indomethacin toxicity in a mouse model. The increased colonic damage in males and higher hepatic toxicity in females caused us to examine the levels of indomethacin and its inactive glucuronide in several murine tissues. We found that male mice showed significantly higher intestinal indomethacin, while the female mice contained significantly higher levels of indomethacin-glucuronide in blood. These data support the conclusion that the trafficking of indomethacin and its metabolites are different in male and female mice and led to greater colonic damage and associated toxicities in males and enhanced splenic and hepatic toxicity in females.

We initially hypothesized that differences in hepatic UDP-glucunosyltransferase (UGT) activities might lead to distinct levels of indomethacin and indomethacin-glucuronide in the intestines and blood of male and female mice. However, we found that there were no differences between male and female mice in the levels of indomethacin and its inactive metabolite indomethacin-glucuronide in the liver. Thus, we next focused on literature evidence for distinct hepatic efflux pathways for indomethacin in males and females. We found that indomethacin-glucuronide, along with other small molecule glucuronides, are subject to efflux from hepatocytes by the multidrug resistance associated protein 2 (MRP2) and MRP3 efflux factors ^25^, members of the ABCC subfamily of membrane transporters. MRP2 is in the apical membrane of hepatocytes and facilitates biliary secretion to the intestines, while MRP3 is localized to the basolateral membrane of hepatocytes and effluxes compounds to the blood (shown in **Figure S3**) ^26^. Furthermore, estrogen has been shown to up-regulate the expression of MRP3 and down-regulates of expression of MRP2 in the female liver (**Figure S3**) ^26^. Thus, indomethacin-glucuronide levels would be expected to be higher in the blood of female animals due to MRP3, while indomethacin-glucuronide would be predominantly trafficked to the bile duct for GI excretion in males due to MRP2. Indomethacin-glucuronide is known to be converted to indomethacin by the gut microbiota ^12^, which would explain the higher measured levels in males and the concomitant higher degree of colonic and associated damage in male animals. By contrast, female mice traffic more indomethacin-glucuronide to the blood, presumably for excretion in the urine, but in doing so appear to subject their spleens and livers to higher degrees of toxicity, as evidenced by the inflammatory cytokine expression outcomes we observed.

The gut microbial β-glucuronidase inhibitor UNC10201652 (GUSi) delivered by oral gavage at 2 mg/kg body weight starting three days before indomethacin significantly alleviated distinct indomethacin-induced toxicities in male and female mice. In males, GUSi blocked colonic immune cell infiltration and *Il-6* expression, and normalized *Tff-3* expression levels in the colon. TFF3 suppresses intestinal inflammation via PAR-2 and TLR/NF-*κ*B signaling ^27^. *Tff-3*^-/-^ mice exhibit impaired mucosal healing, and increased colonic mucosal permeability and inflammation ^28,29^. In our study, we observed notable sex-based differences in *Tff-3* expression in the colon: indomethacin significantly decreased *Tff-3* levels in male mice but caused no change in female mice. Estrogen has been reported to upregulate *Tff-3* expression in breast cancer model ^30,31^, which may explain the higher levels observed in our females. We found that GUSi normalized the *Tff-3* gene expression in colon of male mice treated with indomethacin. GUSi also reduced the translocation of bacteria to the blood in male mice, along with reducing the spleen weight and normalizing the hepatic expression of *Il-6* and *Ppar-γ*. In female mice, GUSi reduced colonic ulcers and *Mcp-1* expression, blocked inflammatory cytokine expression in the liver and spleen, and normalized hepatic *Ppar-α* expression. Furthermore, GUSi blocked the increase in Verrucomicrobia in the female mouse colon. Thus, GUSi alleviates several measures of indomethacin-induced toxicity in this murine model. Still, GUSi used at this dose was insufficient to block all measures of the damage caused by indomethacin. The same GUS inhibitor used at half this dose has been shown to alleviate the toxicity of the consumer product toxin triclosan in mouse model of colitis ^24^, and to prevent the intestinal damage caused by irinotecan in a mouse solid tumor model ^23^. Both triclosan and the active metabolite of irinotecan, SN-38, are sent to the gut as inactive glucuronide conjugates, and these studies support the conclusion that GUSi can block the reactivation of these toxic compounds in the GI tract. For indomethacin, increasing the dose of GUSi and/or the length of time it is employed may be necessary to afford complete protection, something that will be addressed in future studies. Indeed, it was recently demonstrated that a longer-term administration of 2 mg/kg GUSi, 21 days by oral gavage, led to a significant increase in the level of serotonin-glucuronide and decrease the level of active serotonin in the intestines and plasma of male and female mice ^16^.

In terms of gut microbial composition, a published study in male C57BL/6 mice showed that a single oral dose of 10 mg/kg indomethacin alters in six hours the alpha diversity of the fecal microbiota, increasing in the levels of Bacteroidota and decreasing in the levels of Firmicutes ^32^. A second study also showed that oral indomethacin changes the composition of the fecal microbiota within two days. In this case, C57BL/6 mice of unspecified sex that received 10 mg/kg indomethacin orally for two days showed a significant increase in fecal levels of Firmicutes and a concomitant decrease in levels of Bacteroidota ^33^. Furthermore, Yan et al. employed five days of 5mg/kg indomethacin orally in C67BL/6 mice of unspecified sex and found a significant decrease in the relative ratio of Firmicutes to Bacteroidota ^34^. Here, we did not find significant shifts in Firmicutes and Bacteroidota our male and female C57BL/6 mice after five days of oral indomethacin at 5 mg/kg. However, we did observe a significant indomethacin-induced increase in fecal Verrucomicrobia in the female mice, an effect that was blocked by GUSi. It is not immediately clear why females showed this effect. We note that the compositions of the male and female fecal microbiota were significantly different at the start of our study, and it is possible that the male community structure blocked the Verrucomicrobia bloom. In addition, given the reduced levels of indomethacin-induced colonic toxicity in the females, the increased Verrucomicrobia and associated *Akkermansia* genera may be further evidence of a healthier response to indomethacin in the female gut, one that could not be accessed by the males.

Sex is an important biological variable for a range of diseases and therapeutic interventions. NSAIDs are used indiscriminately by individuals of both sexes, even though human males and females contain distinctive gut microbiota profiles, sex hormones, intestinal permeability, and immune responses ^35–40^. There has been growing recognition that disease prevalence, course, and outcomes are significantly different in men and women ^41,42^. Women experience higher rates of adverse drug reactions, generally have a higher percentage of body fat, slower gastric emptying times, and lower glomerular filtration rates than men, all of which can influence NSAID pharmacokinetics ^43^. Here, we define sex-specific toxicities associated with the NSAID indomethacin, as well as evidence for a microbiome-targeted approach to alleviating these injuries. Given the wide use of NSAIDs and the relative lack of understanding of how male and female differ in disease etiologies and interventions, these results may eventually lead to improved human therapeutic outcomes with NSAIDs.

## MATERIALS & METHODS

### Animal experiments

All animal experiments were conducted in accordance with the protocols approved by the Institutional Animal Care and Use Committee of the University of North Carolina at Chapel Hill (IACUC approval number 22-170.0). Mice maintained in a specific pathogen free (SPF) facility with 12-hour light/dark cycle, maintained between 20-23°C with 30-70% relative humidity. Mice had ad libitum access to drinking water and standard chow diet (irradiated Purina PicoLab® Select Rodent 50 IF/6F 5V5R*) for the entire study. At the end of the experiment, the mice were sacrificed by CO2 and cervical dislocation, and the tissues were collected aseptically in a clean operation room located in the SPF facility.

### Animal experiment 1: Effects of indomethacin on the mouse gastrointestinal tract

C57BL/6N (male and female 8-week-old, n=6/group/sex) mice were purchased from Charles River Laboratories and allowed to acclimatize for two weeks prior to study initiation. Mice were orally gavaged with indomethacin (dose = 5mg/kg) or vehicle (Veh; a mixed solvent of 1:9 PEG 400 and PBS) daily for 5 days (see study scheme in **Figure 1A**). Mice were sacrificed 12-18 hours following their last dose of indomethacin or vehicle, and materials and tissues were collected for further analysis.

### Animal experiment 2: Effects of GUS inhibition on indomethacin-treated mice

C57BL/6N (male and female, 8-week-old, n=7/group/sex) mice were purchased from Charles River Laboratories and allowed to acclimatize for two weeks prior to study initiation. Mice were orally gavaged daily with GUS inhibitor (GUSi, UNC10201652, dose = 2 mg/kg) or control (a mixed solvent of 1:9 DMSO and saline) throughout the experiment, as described previously ^24^. After 3 days, mice were treated also with indomethacin (dose = 5mg/kg) or vehicle (Veh, a mixed solvent of 1:9 PEG 400 and PBS) by oral gavage daily for 5 days (see study scheme in **Figure 3A**). GUSi/control were provided in the morning and indomethacin/Veh in the late afternoon. Mice were sacrificed 12-18 hours following their last dose of indomethacin or vehicle, and materials and tissues were collected for further analysis.

### Animal experiment 3: Alteration of gut microbiome on indomethacin-treated mice

C57BL/6N (male and female, 8-week-old, n=7/group/sex) mice were purchased from Charles River Laboratories. At the start of the study, the mice underwent a cage rotation procedure. Fecal samples were collected from all mice. The mice were then randomly grouped and housed, with one mouse per cage for the entire study. Next, the indomethacin treatment and GUSi treatment were administered as described in **Animal experiment 2**. Finally, fecal samples were collected from all mice at the end of the study for further analysis.

### Flow cytometry

Distal colon tissues were dissected, washed with cold PBS, and digested with Hank’s balanced salt solution (HBSS, Lonza) supplemented with 1 mM dithiothreitol (DTT) and 5 mM EDTA at 4 °C (colon epidermal cells) for 4 hours as previously described ^44,45^. The released cells were stained with FITC-conjugated anti-mouse CD45 (BioLegend, Clone: 30-F11), PerCP/Cy5.5-conjugated anti-mouse F4/80 (BioLegend, Clone: BM8), and PE/Cy7-conjugated anti-mouse Ly-6G/Ly-6C (GR-1) (BioLegend, Clone: RB6-8C5) to detect CD45^+^ cells, CD45^+^F4/80^+^ cells and CD45^+^GR-1^+^ cells. Cells were stained with Zombie Violet^TM^ dye (Zombie Violet^TM^ Fixable Viability Kit; BioLegend) according to the manufacturer’s instructions to exclude dead cells. For compensation controls, individual samples of the digested cells were stained separately with each fluorophore. Gating and cell identification strategies were: cell doublets and clumps were eliminated using FSC-A gating and debris was eliminated using FSC-A vs SSC-A. Dead cells were gated out using Zombie Violet™ dye. Flow cytometry data were acquired on a BD LSR Fortessa^TM^ cell analyzer (Becton Dickinson, Franklin Lakes, NJ) and analyzed using FlowJo software (FlowJo, LLC).

### Reverse-Transcriptase-qPCR of inflammatory biomarkers

Total RNA of colon tissues was isolated using Trizol reagent (Ambion) according to the manufacturer’s instructions. RNA was reverse transcribed into cDNA using the High Capacity cDNA Reverse Transcription kit (Applied Biosystem) according to the manufacturer’s instructions. 20 μL PCR reactions were prepared using the Maxima SYBR green Master Mix (Thermo Fisher Scientific), and qPCR was carried out using a DNA Engine Opticon system (Bio-Rad Laboratories). Mouse-specific primer sequences (Integrated DNA Technologies) used to detect inflammatory biomarkers are listed in **Table S1.** The results for the target genes were normalized to *Gapdh* using the 2 ^-ΔΔCT^ method.

### Analysis of *16S rRNA* to quantify bacterial load in tissues and blood

Bacterial load in tissues or blood was analyzed as described previously ^45,46^. Total DNA was extracted from the tissues or blood using QIAamp DNAeasy Blood & Tissue Kit (Qiagen), following the manufacturer’s instructions with the addition of a bead-beating step. The quality of the extracted DNA was measured using a NanoDrop Spectrophotometer (Thermo Fisher Scientific) and qRT-PCR was performed using 5 ng/mL DNA in a DNA Engine Opticon system with Maxima SYBR-Green Master Mix (Applied Biosystems). The sequences of mouse-specific primer (Thermo Fisher Scientific) are listed in Supplementary **Table S1**.

### Histological staining

The dissected colon tissues were fixed in 10% neutral buffered formalin (Thermo Fisher Scientific) for 48 hours. After dehydration, the tissues were embedded in paraffin and sliced (5 μm) by a Rotary Microtome (Leica Biosystems). The slices were dewaxed in serial xylene and rehydrated through ethanol solutions, stained with hematoxylin and eosin (Epredia Richard-Allan Scientific), and images were obtained under light microscope. The ulceration was evaluated by a board-certified veterinary pathologist at the UNC Pathology Core.

### Detection of indomethacin and its metabolites by LC-MS/MS

Mouse tissues/blood were placed in homogenizer tubes with beads and 1mL methanol, then homogenized using a Tissue Lyser II (Qiagen, Valencia, CA). Samples were centrifuged at 10,000 rpm for 3 min. The supernatant was collected and centrifuged again at 14,000 rpm for 5 min, and 500 μL of the supernatant was collected and vacuum centrifuged to dryness. Stable isotope-labeled IND-d4 (Cayman Chemical Company, Ann Arbor, MI) was used as the surrogate standard during the extraction. The extracts were re-dissolved in methanol with the amount that was proportional to sample weights or volumes, then centrifuged (14,000 rpm, 15 min, 4 °C) before the LC-MS/MS analysis for indomethacin (IND) and its metabolite indomethacin-acyl-b-D-glucuronide (IND-G). IND and IND-G were quantified using an Agilent 1200 SL ultrahigh performance liquid chromatography (UHPLC) system coupled with an AB Sciex 4000 Qtrap Mass Spectrometer. Kinetex UPLC C18 column (1.7-μm particles, 2.1 × 50 mm, Phenomenex) was used for chromatographic separation. Data acquisition was performed by multiple reaction monitoring (MRM) in negative ionization mode. The data were analyzed using Analyst software (version 1.6, AB Sciex). For the quantification of IND and IND-G by LC-MS/MS in the different mouse experiments, blank samples from the control group without IND treatment were used as the matrixes for calibration curve standards. During the instrumental analysis, the matrix calibration curve was performed at the beginning and at the end of every sample batch. All reported concentrations were determined based on a standard curve with 7-10 data points.

### DNA extraction from fecal samples

DNA was extracted from mouse fecal samples as described previously ^24,47,48^, using QIAmp DNA Stool Mini Kit (Qiagen, Valencia, CA) following instructions from the manufacturer with an additional bead-beating step. The quantity of the extracted DNA was measured using a NanoDrop Spectrophotometer (Thermo Fisher Scientific). The DNA was then subjected to *16S rRNA* sequencing analysis by Novogene (Sacramento, CA).

### 16S *rRNA* sequencing and analysis

DNA quality was monitored on 1% agarose gels. The V3-V4 hypervariable regions of the bacteria *16S rRNA* gene were amplified with primers 341F (5’-CCTAYGGGRBGCASCAG-3’) and 806R (5’-GGACTACNNGGGTATCTAAT-3’). PCR products were detected on 2% agarose gels by electrophoresis and purified using the Qiagen Gel Extraction Kit (Qiagen, Germany). Sequencing libraries were generated using NEBNext Ultra DNA Library Pre-Kit for Illumina, following manufacturer’s recommendations and index codes were added. The library quality was assessed using the Qubit 2.0 Fluorometer (Thermo Scientific) and Agilent Bioanalyzer 2100 system. The library was sequenced on an Illumina platform and 470 bp paired-end reads were generated.

### Data analysis

All data are expressed as the mean ± standard error of the mean (SEM). For the comparison between treatment groups, Shapiro-Wilk test was used to verify the normality of data. When data were normally distributed, statistical significance was determined using two-side t-test; otherwise, significance was determined by Mann-Whitney test. P values less than 0.05 are reported as statistically significant.

Analysis of differences in abundance patterns among samples was performed by beta diversity using weighted UniFrac distance followed by Principal Coordinate Analysis (PCoA) and non-metric multidimensional scaling (NMDS). The alpha diversity, the richness of the samples, was considered as the number of OTUs present per treatment. We calculated Simpson, Shannon, Chao1 index using QIIME2 version 2019.7.0 and Phytools package 0.7v in R. Statistical tests were performed in R. Kruskal-Wallis rank sum test was used to compare alpha diversity indexes among treatments. The beta diversity was calculated using Phytools 0.7v. The weighted UniFrac distance method was used to create the matrix followed by PCoA to compare similarity among treatments. All plots were obtained using ggplot2.

## ACKNOWLEDGMENTS

This research was supported by NIH R35 award GM152079 (to M.R.R.). The histological staining was supported by NIH P30 DK034987 (Center for Gastrointestinal Biology and Disease Histology Core) and conducted in the memory of Research Specialist Carolyn B. Suitt.

## AUTHOR CONTRIBUTIONS

J.Z. performed the animal experiments. J.Z and J.J.S. analyzed the data. J.Y. performed the LC/MS/MS experiment. R.S.S. quantified the ulcer number in intestines. J.Z. and M.R.R. wrote the original manuscript. J.Z., A.P.B. and M.R.R. revised the manuscript. J.Z. and M.R.R. designed the study.

## DATA AVAILABILITY STATEMENT

All data generated or analyzed during this study are included in this published article (and its Supplementary Information files).

## APPROVAL FOR ANIMAL EXPERIMENTS AND ETHICS DECLARATIONS

This study was designed and conducted in accordance with the ARRIVE 2.0 guidelines for reporting animal research. All animals received appropriate anesthesia and analgesia to minimize suffering, and humane endpoints were established prior to the study and followed the animal protocols approved by the Institutional Animal Care and Use Committee of the University of North Carolina at Chapel Hill (IACUC approval number 22-170.0). A full checklist document “The ARRIVE Essential 10” is attached to the Supporting Information.

## ADDITIONAL INFORMATION

Supplementary Information is attached.

## REFERENCES

1 Day, R. O. & Graham, G. G. Non-steroidal anti-inflammatory drugs (NSAIDs). Bmj 346 (2013).

2 Serhan, C. N. Pro-resolving lipid mediators are leads for resolution physiology. Nature 510, 92–101 (2014).

3 Boelsterli, U. A., Redinbo, M. R. & Saitta, K. S. Multiple NSAID-induced hits injure the small intestine: underlying mechanisms and novel strategies. toxicological sciences 131, 654–667 (2013).

4 Crittenden, S. et al. Prostaglandin E2 promotes intestinal inflammation via inhibiting microbiota-dependent regulatory T cells. Science advances 7, eabd7954 (2021).

5 Wilcox, C. M., Alexander, L. N., Cotsonis, G. A. & Clark, W. S. Nonsteroidal antiinflammatory drugs are associated with both upper and lower gastrointestinal bleeding. Digestive diseases and sciences 42, 990–997 (1997).

6 Wolfe, M. M., Lichtenstein, D. R. & Singh, G. Gastrointestinal toxicity of nonsteroidal antiinflammatory drugs. New England Journal of Medicine 340, 1888–1899 (1999).

7 Uejima, M., Kinouchi, T., Kataoka, K., Hiraoka, I. & Ohnishi, Y. Role of intestinal bacteria in ileal ulcer formation in rats treated with a nonsteroidal antiinflammatory drug. Microbiology and immunology 40, 553–560 (1996).

8 Syer, S. D. et al. NSAID enteropathy and bacteria: a complicated relationship. Journal of gastroenterology 50, 387–393 (2015).

9 Maseda, D. & Ricciotti, E. NSAID–gut microbiota interactions. Frontiers in pharmacology 11, 1153 (2020).

10 Biernat, K. A. et al. Structure, function, and inhibition of drug reactivating human gut microbial β-glucuronidases. Scientific reports 9, 1–15 (2019).

11 LoGuidice, A., Wallace, B. D., Bendel, L., Redinbo, M. R. & Boelsterli, U. A. Pharmacologic targeting of bacterial β-glucuronidase alleviates nonsteroidal anti-inflammatory drug-induced enteropathy in mice. Journal of Pharmacology and Experimental Therapeutics 341, 447–454 (2012).

12 Saitta, K. S. et al. Bacterial β-glucuronidase inhibition protects mice against enteropathy induced by indomethacin, ketoprofen or diclofenac: mode of action and pharmacokinetics. Xenobiotica 44, 28–35 (2014).

13 Yauw, S. T. et al. Microbial glucuronidase inhibition reduces severity of diclofenac-induced anastomotic leak in rats. Surgical Infections 19, 417–423 (2018).

14 Kwa, M., Plottel, C. S., Blaser, M. J. & Adams, S. The intestinal microbiome and estrogen receptor–positive female breast cancer. Journal of the National Cancer Institute 108, djw029 (2016).

15 Ervin, S. M. et al. Gut microbial β-glucuronidases reactivate estrogens as components of the estrobolome that reactivate estrogens. Journal of Biological Chemistry 294, 18586–18599 (2019).

16 Simpson, J. B. et al. Gut microbial β-glucuronidases influence endobiotic homeostasis and are modulated by diverse therapeutics. Cell Host & Microbe 32, 925–944. e910 (2024).

17 Crofford, L. J. Use of NSAIDs in treating patients with arthritis. Arthritis research & therapy 15, 1–10 (2013).

18 Favalli, E. G. et al. Sex and management of rheumatoid arthritis. Clinical reviews in allergy & immunology 56, 333–345 (2019).

19 Fosbøl, E. L. et al. The pattern of use of non−steroidal anti−inflammatory drugs (NSAIDs) from 1997 to 2005: a nationwide study on 4.6 million people. Pharmacoepidemiology and drug safety 17, 822–833 (2008).

20 Edogawa, S. et al. Sex differences in NSAID-induced perturbation of human intestinal barrier function and microbiota. The FASEB Journal 32, 6615 (2018).

21 Chen, C. et al. The roles of estrogen and estrogen receptors in gastrointestinal disease. Oncology letters 18, 5673–5680 (2019).

22 Palileo, C. & Kaunitz, J. D. Gastrointestinal defense mechanisms. Current opinion in gastroenterology 27, 543–548 (2011).

23 Bhatt, A. P. et al. Targeted inhibition of gut bacterial β-glucuronidase activity enhances anticancer drug efficacy. Proceedings of the National Academy of Sciences 117, 7374–7381 (2020).

24 Zhang, J. et al. Microbial enzymes induce colitis by reactivating triclosan in the mouse gastrointestinal tract. Nature communications 13, 1–14 (2022).

25 Kouzuki, H., Suzuki, H. & Sugiyama, Y. Pharmacokinetic study of the hepatobiliary transport of indomethacin. Pharmaceutical research 17, 432–438 (2000).

26 Mineiro, R. et al. Regulation of ABC transporters by sex steroids may explain differences in drug resistance between sexes. Journal of physiology and biochemistry 79, 467–487 (2023).

27 Yang, Y., Lin, Z., Lin, Q., Bei, W. & Guo, J. Pathological and therapeutic roles of bioactive peptide trefoil factor 3 in diverse diseases: recent progress and perspective. Cell death & disease 13, 62 (2022).

28 Mashimo, H., Wu, D.-C., Podolsky, D. K. & Fishman, M. C. Impaired defense of intestinal mucosa in mice lacking intestinal trefoil factor. Science 274, 262–265 (1996).

29 Podolsky, D. K., Gerken, G., Eyking, A. & Cario, E. Colitis-associated variant of TLR2 causes impaired mucosal repair because of TFF3 deficiency. Gastroenterology 137, 209–220 (2009).

30 Chen, S. et al. TFF3 facilitates dormancy of anti-estrogen treated ER+ mammary carcinoma. Communications Medicine 5, 45 (2025).

31 Clemons, M. & Goss, P. Estrogen and the risk of breast cancer. New England Journal of Medicine 344, 276–285 (2001).

32 Liang, X. et al. Bidirectional interactions between indomethacin and the murine intestinal microbiota. Elife 4, e08973 (2015).

33 Xiao, X. et al. Gut microbiota mediates protection against enteropathy induced by indomethacin. Scientific reports 7, 40317 (2017).

34 Yan, Y., Zhou, X., Guo, K., Zhou, F. & Yang, H. Chlorogenic acid protects against indomethacin-induced inflammation and mucosa damage by decreasing bacteroides-derived LPS. Frontiers in immunology 11, 1125 (2020).

35 Kim, N. Sex difference of gut microbiota. Sex/gender-specific medicine in the gastrointestinal diseases, 363–377 (2022).

36 Yoon, K. & Kim, N. Roles of sex hormones and gender in the gut microbiota. Journal of neurogastroenterology and motility 27, 314 (2021).

37 Ahnstedt, H. et al. Sex differences in T cell immune responses, gut permeability and outcome after ischemic stroke in aged mice. Brain, behavior, and immunity 87, 556–567 (2020).

38 Meleine, M. & Matricon, J. Gender-related differences in irritable bowel syndrome: potential mechanisms of sex hormones. World Journal of Gastroenterology: WJG 20, 6725 (2014).

39 Kim, Y. S. & Kim, N. Sex-gender differences in irritable bowel syndrome. Journal of neurogastroenterology and motility 24, 544 (2018).

40 Homma, H. et al. The female intestine is more resistant than the male intestine to gut injury and inflammation when subjected to conditions associated with shock states. American Journal of Physiology-Gastrointestinal and Liver Physiology 288, G466–G472 (2005).

41 Farkouh, A. et al. Sex-related differences in drugs with anti-inflammatory properties. Journal of clinical medicine 10, 1441 (2021).

42 Bernstein, S. R., Kelleher, C. & Khalil, R. A. Gender-based research underscores sex differences in biological processes, clinical disorders and pharmacological interventions. Biochemical Pharmacology, 115737 (2023).

43 Zucker, I. & Prendergast, B. J. Sex differences in pharmacokinetics predict adverse drug reactions in women. Biology of sex differences 11, 1–14 (2020).

44 Wang, W. et al. Lipidomic profiling reveals soluble epoxide hydrolase as a therapeutic target of obesity-induced colonic inflammation. Proceedings of the National Academy of Sciences 115, 5283–5288 (2018).

45 Zhang, J. et al. Thermally processed oil exaggerates colonic inflammation and colitis-associated colon tumorigenesis in mice. Cancer Prevention Research 12, 741–750 (2019).

46 Lei, L., Yang, J., Zhang, J. & Zhang, G. The lipid peroxidation product EKODE exacerbates colonic inflammation and colon tumorigenesis. Redox biology 42, 101880 (2021).

47 Zhang, J. et al. CYP eicosanoid pathway mediates colon cancer−promoting effects of dietary linoleic acid. The FASEB Journal 37, e23009 (2023).

48 Wang, W. et al. Oxidized polyunsaturated fatty acid promotes colitis and colitis-associated tumorigenesis in mice. Journal of Crohn’s and Colitis, jjae148 (2024).

