## Supplementary Information for "Sex-Dependent Responses in Mice to Indomethacin-Induced Injury and Gut Microbiome-Targeted Alleviation"

**Table S1. Sequences of primers in qRT-PCR**

| Gene | Forward primer | Reverse primer |
|---|---|---|
| *Gapdh* | AGGTCGGTGTGAACGGATTTG | TGTAGACCATGTAGTTGAGGTCA |
| *Tnf-α* | CCCTCACACTCAGATCATCTTCT | GCTACGACGTGGGCTACAG |
| *Il-6* | TAGTCCTTCCTACCCCAATTTCC | TTGGTCCTTAGCCACTCCTTC |
| *Il-1β* | GCAACTGTTCCTGAACTCAACT | ATCTTTTGGGGTCCGTCAACT |
| *Mcp-1* | TTAAAAACCTGGATCGGAACCAA | GCATTAGCTTCAGATTTACGGGT |
| *Zo-1* | GGGGCCTACACTGATCAAGA | TGGAGATGAGGCTTCTGCTT |
| *Tff3* | CCTGGTTGCTGGGTCCTCTG | GCCACGGTTGTTACACTGCTC |
| *Muc2* | GCTGACGAGTGGTTGGTGAATG | GATGAGGTGGCAGACAGGAGAC |
| *Cd36* | GGAGCCATCTTTGAGCCTTCA | GAACCAAACTGAGGAATGGATCT |
| *Ppar-α* | AGAGCCCCATCTGTCCTCTC | ACTGGTAGTCTGCAAAACCAAA |
| *Ppar-γ* | TTTTCCGAAGAACCATCCGATT | ATGGCATTGTGAGACATCCCC |
| *16srRNA* | TCGTCGGCAGCGTCAGATGTGTATAA GAGACAGCCTACGGGNGGCWGCAG | GTCTCGTGGGCTCGGAGATGTGTATA AGAGACAGGACTACHVGGGTATCTAATCC |
| **Primers for sequencing** | **Forward primer (341F)** | **Reverse primer (806R)** |
| 16S rRNA | CCTAYGGGRBGCASCAG | GGACTACNNGGGTATCTAAT |

**Table S2. Comparison of the phylum-level differences in abundance of fecal microbiota by sex factor and by treatment factor**. P<0.05. F: Female; M: Male; IND: indomethacin.

| Taxonomy to Phylum | Group | Estimate | P value |
|---|---|---|---|
| Bacteroidota | F - M | 22.8577 | <0.0001 |
| Firmicutes | F - M | -24.8679 | <0.0001 |
| Deferribacterota | F - M | -0.0025 | 0.0229 |
| **Taxonomy to Phylum** | **Group** | **Estimate** | **P value** |
| Verrucomicrobiota | Ctrl - Ctrl+IND | -7.2871 | 0.0083 |
| Verrucomicrobiota | Ctrl+IND - GUSi+IND | 8.6417 | 0.0017 |
| Bacteroidota | GUSi - GUSi+IND | 11.0826 | 0.0242 |
| Firmicutes | GUSi - GUSi+IND | -13.4042 | 0.0259 |

**Table S3. Comparison of the order-level differences in abundance of fecal microbiota by sex factor and by treatment factor**. We noticed a significant difference in the abundance of Verrucomicrobiota at the phylum level in the IND- or GUSi-treated groups; the taxa of order-level are highlighted in this table. P<0.05. F: Female; M: Male; IND: indomethacin.

| Taxonomy to Order | Group | Estimate | P value |
|---|---|---|---|
| Firmicutes__Clostridia__Oscillospirales | F - M | -7.3288 | <0.0001 |
| Bacteroidota__Bacteroidia__Bacteroidales | F - M | 23.9169 | <0.0001 |
| Firmicutes__Clostridia__Eubacteriales | F - M | -0.0118 | <0.0001 |
| Bacteroidota__Bacteroidia__Flavobacteriales | F - M | -1.0590 | <0.0001 |
| Firmicutes__Clostridia__Peptococcales | F - M | -0.0319 | <0.0001 |
| Firmicutes__Clostridia__Christensenellales | F - M | -0.2271 | <0.0001 |
| Firmicutes__Incertae_Sedis__DTU014 | F - M | -0.0065 | <0.0001 |
| Firmicutes__Clostridia__Clostridia | F - M | -0.0250 | <0.0001 |
| Firmicutes__Clostridia__Lachnospirales | F - M | -18.0459 | <0.0001 |
| Firmicutes__Bacilli__Erysipelotrichales | F - M | 0.4355 | 0.0002 |

| | | | |
|---|---|---|---|
| Firmicutes__Clostridia__Monoglobales | F - M | -0.0133 | 0.0043 |
| Proteobacteria__Gammaproteobacteria__Enterobacterales | F - M | 0.0062 | 0.0065 |
| Deferribacterota__Deferribacteres__Deferribacterales | F - M | -0.0025 | 0.0229 |
| Proteobacteria__Alphaproteobacteria__Rhodospirillales | F - M | -0.0211 | 0.0242 |
| Actinobacteriota__Coriobacteriia__Coriobacteriales | F - M | -0.2050 | 0.0265 |
| Verrucomicrobiota__Lentisphaeria__Victivallales | F - M | -0.0012 | 0.0328 |
| Firmicutes__Clostridia__Clostridiales | F - M | -0.9119 | 0.0442 |
| **Taxonomy to Order** | **Group** | **Estimate** | **P value** |
| Proteobacteria__Gammaproteobacteria__Enterobacterales | Ctrl - Ctrl+IND | -0.0122 | 0.0017 |
| Verrucomicrobiota__Verrucomicrobiae__Verrucomicrobiales | Ctrl - Ctrl+IND | -7.2870 | 0.0083 |
| Firmicutes__Clostridia__Eubacteriales | Ctrl - GUSi+IND | -0.0065 | 0.0200 |
| Proteobacteria__Gammaproteobacteria__Enterobacterales | Ctrl+IND - GUSi+IND | 0.0121 | 0.0008 |
| Verrucomicrobiota__Verrucomicrobiae__Verrucomicrobiales | Ctrl+IND - GUSi+IND | 8.6421 | 0.0017 |
| Firmicutes__Clostridia__Eubacteriales | Ctrl+IND - GUSi+IND | -0.0063 | 0.0268 |
| Firmicutes__Clostridia__Clostridia | GUSi - GUSi+IND | 0.0222 | 0.0227 |
| Firmicutes__Incertae_Sedis__DTU014 | GUSi - GUSi+IND | 0.0049 | 0.0275 |
| Bacteroidota__Bacteroidia__Bacteroidales | GUSi - GUSi+IND | 10.5506 | 0.0308 |
| Firmicutes__Clostridia__Lachnospirales | GUSi - GUSi+IND | -12.9233 | 0.0373 |

**Table S4. Comparison of the genus-level differences in abundance of fecal microbiota by sex factor and by treatment factor**. We noticed a significant difference in the abundance of Verrucomicrobiota at the phylum level in the IND- or GUSi-treated groups; the taxa of genus-level are highlighted in this table. P<0.05. F: Female; M: Male; IND: indomethacin.

| Taxonomy to Genus | Group | Estimate | P value |
|---|---|---|---|
| Bacteroidota__Bacteroidia__Bacteroidales__Muribaculaceae__Muribaculaceae | F - M | 43.2965 | <0.0001 |
| Firmicutes__Clostridia__Christensenellales__Christensenellaceae__uncultured | F - M | -0.0443 | <0.0001 |
| Firmicutes__Clostridia__Oscillospirales__Oscillospiraceae__NK4A214 group | F - M | -0.0936 | <0.0001 |
| Bacteroidota__Bacteroidia__Bacteroidales__Bacteroidaceae__Bacteroides | F - M | -13.6430 | <0.0001 |
| Firmicutes__Clostridia__Peptostreptococcales Tissierellales__Anaerovoracaceae__Family XIII UCG.001 | F - M | -0.0675 | <0.0001 |
| Firmicutes__Clostridia__Eubacteriales__Anaerofustaceae__Anaerofustis | F - M | -0.0118 | <0.0001 |
| Firmicutes__Clostridia__Peptostreptococcales Tissierellales__Anaerovoracaceae__Family XIII AD3011 group | F - M | -0.0232 | <0.0001 |
| Bacteroidota__Bacteroidia__Flavobacteriales__Flavobacteriaceae__uncultured | F - M | -1.0591 | <0.0001 |
| Bacteroidota__Bacteroidia__Bacteroidales__Marinifilaceae__Butyricimonas | F - M | -5.5649 | <0.0001 |
| Firmicutes__Clostridia__Peptococcales__Peptococcaceae__uncultured | F - M | -0.0319 | <0.0001 |
| Firmicutes__Clostridia__Lachnospirales__Lachnospiraceae__Roseburia | F - M | -1.5374 | <0.0001 |
| Firmicutes__Clostridia__Lachnospirales__Defluviitaleaceae__Defluviitaleaceae UCG.011 | F - M | -0.0632 | <0.0001 |
| Firmicutes__Clostridia__Oscillospirales__Oscillospiraceae__UCG.005 | F - M | -0.6438 | <0.0001 |
| Firmicutes__Clostridia__Oscillospirales__Oscillospiraceae__uncultured | F - M | -1.7762 | <0.0001 |
| Firmicutes__Clostridia__Lachnospirales__Lachnospiraceae__Acetatifactor | F - M | -0.1408 | <0.0001 |
| Firmicutes__Incertae_Sedis__DTU014__DTU014__DTU014 | F - M | -0.0065 | <0.0001 |
| Firmicutes__Clostridia__Oscillospirales__Butyricicoccaceae__UCG.009 | F - M | -0.0784 | <0.0001 |
| Firmicutes__Clostridia__Lachnospirales__Lachnospiraceae__Eubacterium ventriosum group | F - M | 0.4628 | <0.0001 |
| Firmicutes__Clostridia__Peptostreptococcales Tissierellales__Anaerovoracaceae__Eubacterium_nodatum_group | F - M | 0.0525 | <0.0001 |
| Firmicutes__Clostridia__Lachnospirales__Lachnospiraceae__Lachnospiraceae FCS020 group | F - M | -0.1137 | <0.0001 |
| Firmicutes__Clostridia__Oscillospirales__Ruminococcaceae__UBA1819 | F - M | 0.0155 | <0.0001 |
| Firmicutes__Clostridia__Christensenellales__Christensenellaceae__Christensenellaceae R.7 group | F - M | -0.1827 | <0.0001 |
| Firmicutes__Clostridia__Oscillospirales__Ruminococcaceae__Candidatus Soleaferrea | F - M | -0.0045 | 0.0001 |

| | | | |
|---|---|---|---|
| Firmicutes__Clostridia__Peptostreptococcales Tissierellales__Anaerovoracaceae__Anaerovorax | F - M | -0.0058 | 0.0001 |
| Firmicutes__Bacilli__Erysipelotrichales__Erysipelatoclostridiaceae__Erysipelatoclostridium | F - M | 0.4519 | 0.0001 |
| Firmicutes__Clostridia__Oscillospirales__Ruminococcaceae__Eubacterium siraeum group | F - M | -0.0865 | 0.0001 |
| Firmicutes__Clostridia__Oscillospirales__Oscillospiraceae__Colidextribacter | F - M | -0.5755 | 0.0002 |
| Firmicutes__Clostridia__Lachnospirales__Lachnospiraceae__Lachnospiraceae NK4A136 group | F - M | -7.4580 | 0.0002 |
| Firmicutes__Bacilli__Erysipelotrichales__Erysipelatoclostridiaceae__Erysipelatoclostridiaceae | F - M | -0.0071 | 0.0003 |
| Firmicutes__Clostridia__Lachnospirales__Lachnospiraceae__uncultured | F - M | -0.6137 | 0.0004 |
| Firmicutes__Clostridia__Oscillospirales__UCG.010__UCG.010 | F - M | -0.1164 | 0.0004 |
| Firmicutes__Clostridia__Oscillospirales__Ruminococcaceae__Harryflintia | F - M | -0.0097 | 0.0006 |
| Firmicutes__Clostridia__Oscillospirales__Oscillospiraceae__Oscillibacter | F - M | -0.5584 | 0.0010 |
| Firmicutes__Clostridia__Oscillospirales__Ruminococcaceae__Negativibacillus | F - M | -0.0096 | 0.0014 |
| Firmicutes__Clostridia__Lachnospirales__Lachnospiraceae__ASF356 | F - M | -0.1032 | 0.0015 |
| Firmicutes__Bacilli__Erysipelotrichales__Erysipelotrichaceae__Clostridium innocuum group | F - M | 0.0115 | 0.0015 |
| Actinobacteriota__Coriobacteriia__Coriobacteriales__Eggerthellaceae__Gordonibacter | F - M | -0.0132 | 0.0018 |
| Firmicutes__Clostridia__Oscillospirales__Clostridium methylpentosum group__Clostridium methylpentosum group | F - M | -0.0092 | 0.0020 |
| Firmicutes__Clostridia__Lachnospirales__Lachnospiraceae__Tyzzerella | F - M | -0.0035 | 0.0022 |
| Firmicutes__Clostridia__Oscillospirales__Oscillospiraceae__UCG.003 | F - M | -0.0728 | 0.0031 |
| Firmicutes__Clostridia__Oscillospirales__Ruminococcaceae__Ruminococcus | F - M | -1.7124 | 0.0035 |
| Firmicutes__Clostridia__Lachnospirales__Lachnospiraceae__Lachnospiraceae_NK4B4_group | F - M | -0.0449 | 0.0038 |
| Firmicutes__Clostridia__Monoglobales__Monoglobaceae__Monoglobus | F - M | -0.0133 | 0.0043 |
| Firmicutes__Clostridia__Peptostreptococcales Tissierellales__Anaerovoracaceae__Eubacterium brachy group | F - M | -0.0201 | 0.0071 |
| Firmicutes__Bacilli__Erysipelotrichales__Erysipelotrichaceae__Dubosiella | F - M | 0.0419 | 0.0085 |
| Firmicutes__Clostridia__Oscillospirales__Ruminococcaceae__uncultured | F - M | -0.1238 | 0.0105 |
| Firmicutes__Clostridia__Lachnospirales__Lachnospiraceae__Robinsoniella | F - M | 0.0052 | 0.0141 |
| Firmicutes__Clostridia__Oscillospirales__Ruminococcaceae__Paludicola | F - M | -0.0040 | 0.0145 |
| Firmicutes__Clostridia__Oscillospirales__Ruminococcaceae__Incertae Sedis | F - M | -0.1633 | 0.0148 |
| Firmicutes__Bacilli__Erysipelotrichales__Erysipelotrichaceae__Erysipelotrichaceae | F - M | -0.0121 | 0.0160 |

| | | | |
|---|---|---|---|
| Firmicutes__Clostridia__Lachnospirales__Lachnospiraceae__Blautia | F - M | 1.2651 | 0.0162 |
| Firmicutes__Clostridia__Oscillospirales__Oscillospiraceae__UCG.007 | F - M | -0.0026 | 0.0175 |
| Deferribacterota__Deferribacteres__Deferribacterales__Deferribacteraceae__Mucispirillum | F - M | -0.0025 | 0.0229 |
| Proteobacteria__Alphaproteobacteria__Rhodospirillales__uncultured__uncultured | F - M | -0.0211 | 0.0242 |
| Actinobacteriota__Coriobacteriia__Coriobacteriales__Coriobacteriales_Incertae Sedis uncultured | F - M | -0.0006 | 0.0249 |
| Firmicutes__Clostridia__Clostridia__Hungateiclostridiaceae__Ruminiclostridium | F - M | -0.0045 | 0.0252 |
| Verrucomicrobiota__Lentisphaeria__Victivallales__Victivallaceae__Victivallis | F - M | -0.0012 | 0.0328 |
| Actinobacteriota__Coriobacteriia__Coriobacteriales__Eggerthellaceae__Adlercreutzia | F - M | -0.1036 | 0.0411 |
| Firmicutes__Clostridia__Oscillospirales__Ruminococcaceae__Fournierella | F - M | -0.0005 | 0.0455 |
| Firmicutes__Bacilli__Erysipelotrichales__Erysipelotrichaceae__uncultured | F - M | -0.0577 | 0.0462 |

| Taxonomy to Genus | Group | Estimate | P value |
|---|---|---|---|
| Verrucomicrobiota__Verrucomicrobiae__Verrucomicrobiales__Akkermansiaceae__Akkermansia | Ctrl - Ctrl+IND | -7.2870 | 0.0083 |
| Firmicutes__Clostridia__Oscillospirales__Oscillospiraceae__Colidextribacter | Ctrl - Ctrl+IND | -0.4845 | 0.0375 |
| Firmicutes__Clostridia__Christensenellales__Christensenellaceae__uncultured | Ctrl - Ctrl+IND | 0.0142 | 0.0424 |
| Firmicutes__Clostridia__Lachnospirales__Lachnospiraceae__Lachnospiraceae_FCS020_group | Ctrl - Ctrl+IND | -0.0741 | 0.0474 |
| Bacteroidota__Bacteroidia__Bacteroidales__Muribaculaceae__Muribaculaceae | Ctrl - GUSi+IND | 9.5748 | 0.0148 |
| Firmicutes__Clostridia__Eubacteriales__Anaerofustaceae__Anaerofustis | Ctrl - GUSi+IND | -0.0065 | 0.0200 |
| Verrucomicrobiota__Verrucomicrobiae__Verrucomicrobiales__Akkermansiaceae__Akkermansia | Ctrl+IND - GUSi+IND | 8.6421 | 0.0017 |
| Firmicutes__Clostridia__Eubacteriales__Anaerofustaceae__Anaerofustis | Ctrl+IND - GUSi+IND | -0.0063 | 0.0268 |
| Firmicutes__Clostridia__Oscillospirales__UCG.010__UCG.010 | GUSi - GUSi+IND | 0.1666 | 0.0018 |
| Firmicutes__Clostridia__Oscillospirales__Oscillospiraceae__Oscillibacter | GUSi - GUSi+IND | -0.5693 | 0.0137 |
| Firmicutes__Clostridia__Oscillospirales__Oscillospiraceae__Colidextribacter | GUSi - GUSi+IND | -0.5421 | 0.0171 |
| Firmicutes__Incertae_Sedis__DTU014__DTU014__DTU014 | GUSi - GUSi+IND | 0.0049 | 0.0275 |
| Firmicutes__Clostridia__Lachnospirales__Lachnospiraceae__Tuzzerella | GUSi - GUSi+IND | -0.1300 | 0.0373 |
| Firmicutes__Clostridia__Peptostreptococcales Tissierellales__Anaerovoracaceae__Family_XIII_UCG.001 | GUSi - GUSi+IND | -0.0253 | 0.0391 |
| Firmicutes__Clostridia__Oscillospirales__Ruminococcaceae__uncultured | GUSi - GUSi+IND | -0.1742 | 0.0491 |

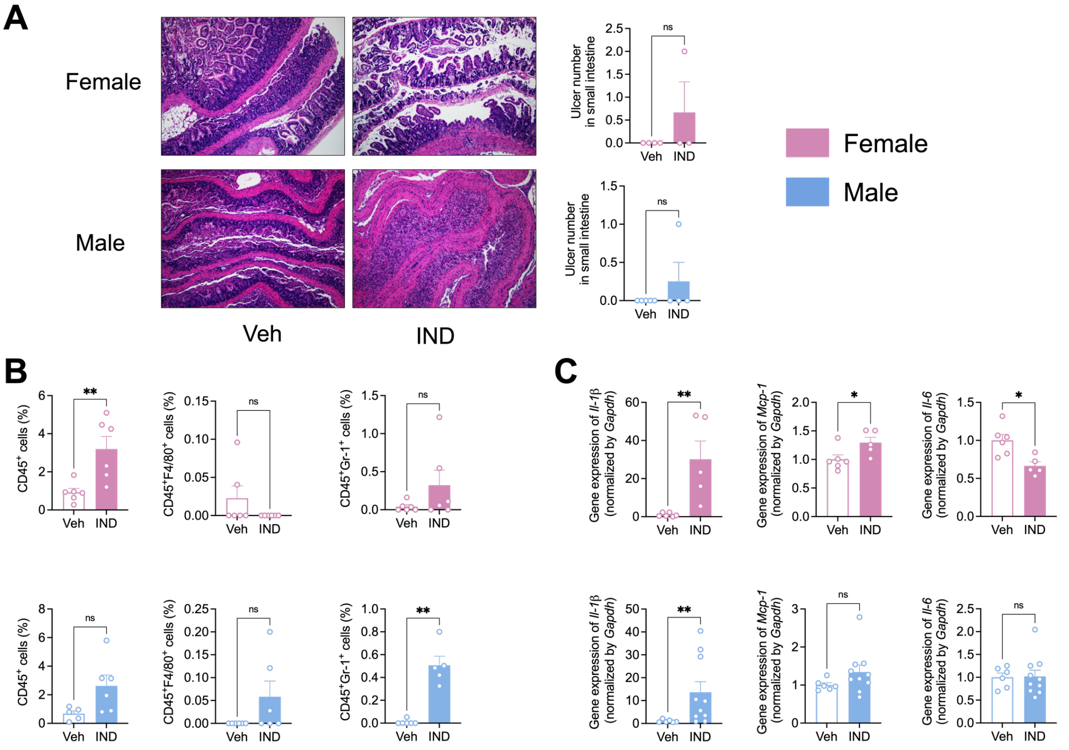

**Figure S1. The effect of indomethacin in small intestine (ileum) in both sexes of mice. A.** Indomethacin induced ulceration in small intestine. Left: representative H&E histological images of small intestine. Right: ulcer numbers in small intestine. **B.** Immune cell infiltration in small intestine: $CD45^+$ cells, $CD45^+F4/80^+$ cells and $CD45^+Gr\text{-}1^+$ cells. **C.** Gene expression of pro-inflammatory cytokines in small intestine. The data are mean ± SEM. *P < 0.05, **P < 0.01, ns. No significance.

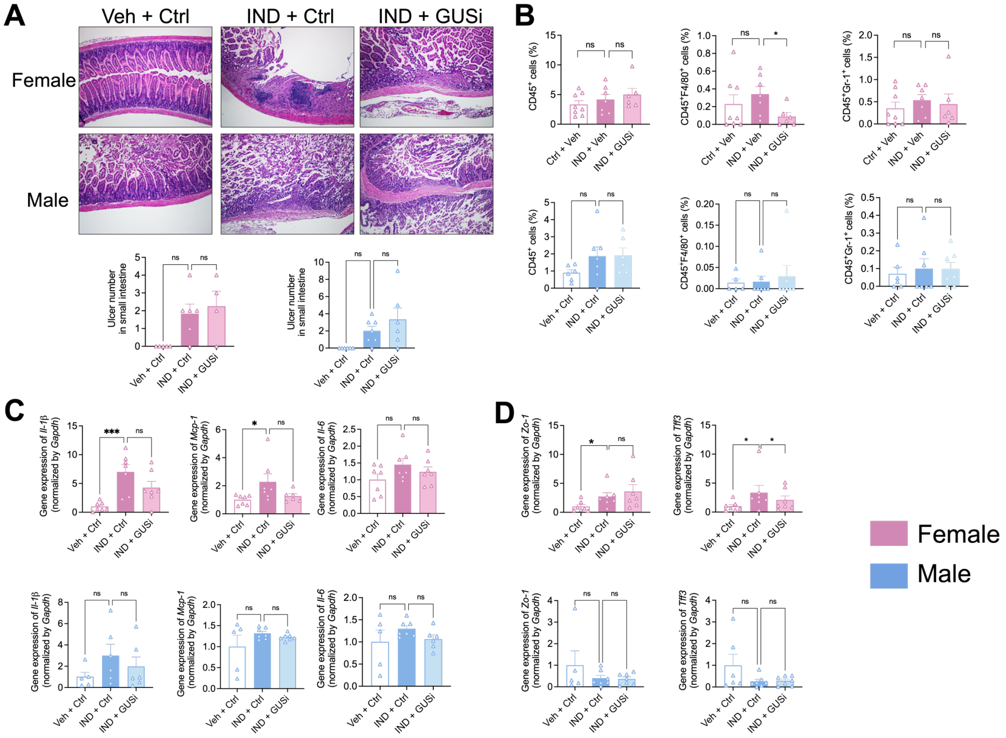

**Figure S2. The effect of inhibition microbial β-glucuronidase (β-GUS) enzymes in indomethacin-induced small intestine. A.** Ulcer number in small intestine. Left: representative H&E histological images of small intestine. Right: ulcer numbers in small intestine. **B.** Gene expression of intestinal leakage markers in small intestine. **C.** Gene expression of pro-inflammatory cytokines in small intestine. **D**. Gene expression of intestinal permeability markers in small intestine. The data are mean ± SEM. *P < 0.05, **P < 0.01, ***P < 0.001. ns. No significance.

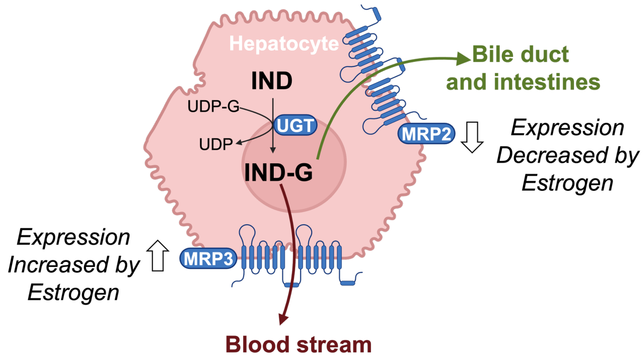

**Figure S3. Transport of indomethacin-glucuronide by MRP2 and MRP3 in hepatocytes.** Indomethacin is metabolized by hepatic UDP-glucuronosyltransferase (UGT) to form indomethacin-glucuronide, which is then transported either via MRP2 to the bile duct and intestines or via MRP3 to the blood stream. Estrogen significantly influences MRP2 and MRP3 expression, potentially creating sex-dependent differences in drug disposition. ( Figure created with BioRender.com).

#### Approval for Animal Experiments and Ethics Declarations

This study was designed and conducted in accordance with the ARRIVE 2.0 guidelines for reporting animal research. All experimental protocols involving animals were approved Institutional Animal Care and Use Committee of the University of North Carolina at Chapel Hill (IACUC approval number 22-170.0). Sample size was determined using a priori power analysis conducted with G*Power 3.1 software. Based on previous similar study (LoGuidice et al. 2012) we calculated the minimum sample size required using the following parameters: two-tailed t-test, significance level $\alpha = 0.05$, statistical power $(1-\beta) = 0.8$, and expected effect size $d = 0.85$. This analysis indicated that a minimum of $n = 6$ animals per group would be necessary to detect the expected effect. To account for potential experimental attrition of approximately 10%, the final sample size was established at n = 6-7 animals per experimental group. Animals were randomly allocated to experimental groups using a body weight-stratified randomization approach. This stratified randomization method minimized potential confounding effects of baseline body weight differences between vehicle group and treatment groups while maintaining the benefits of randomization. All animals received appropriate anesthesia and analgesia to minimize suffering, and humane endpoints were established prior to the study and followed the animal protocols approved by IACUC. Complete details of animal husbandry, housing conditions, and welfare monitoring are provided in the methods section.

1 LoGuidice, A., Wallace, B. D., Bendel, L., Redinbo, M. R. & Boelsterli, U. A. Pharmacologic targeting of bacterial β-glucuronidase alleviates nonsteroidal anti-inflammatory drug-induced enteropathy in mice. *Journal of Pharmacology and Experimental Therapeutics* **341**, 447-454 (2012).

NOTE: Please save this file locally before filling in the table, DO NOT work on the file within your internet browser as changes will not be saved. Adobe Acrobat Reader (available free here) is recommended for completion.

ARRIVE

### The ARRIVE guidelines 2.0: author checklist

#### The ARRIVE Essential 10

These items are the basic minimum to include in a manuscript. Without this information, readers and reviewers cannot assess the reliability of the findings.

| Item | | Recommendation | Section/line number, or reason for not reporting |
|---|---|---|---|
| **Study design** | 1 | For each experiment, provide brief details of study design including: | |
| | | a. The groups being compared, including control groups. If no control group has been used, the rationale should be stated. | Yes |
| | | b. The experimental unit (e.g. a single animal, litter, or cage of animals). | Yes |
| **Sample size** | 2 | a. Specify the exact number of experimental units allocated to each group, and the total number in each experiment. Also indicate the total number of animals used. | Yes |
| | | b. Explain how the sample size was decided. Provide details of any *a priori* sample size calculation, if done. | Yes |
| **Inclusion and exclusion criteria** | 3 | a. Describe any criteria used for including and excluding animals (or experimental units) during the experiment, and data points during the analysis. Specify if these criteria were established *a priori*. If no criteria were set, state this explicitly. | Yes |
| | | b. For each experimental group, report any animals, experimental units or data points not included in the analysis and explain why. If there were no exclusions, state so. | Yes |
| | | c. For each analysis, report the exact value of *n* in each experimental group. | Yes |
| **Randomisation** | 4 | a. State whether randomisation was used to allocate experimental units to control and treatment groups. If done, provide the method used to generate the randomisation sequence. | Yes |
| | | b. Describe the strategy used to minimise potential confounders such as the order of treatments and measurements, or animal/cage location. If confounders were not controlled, state this explicitly. | Yes |
| **Blinding** | 5 | Describe who was aware of the group allocation at the different stages of the experiment (during the allocation, the conduct of the experiment, the outcome assessment, and the data analysis). | Yes |
| **Outcome measures** | 6 | a. Clearly define all outcome measures assessed (e.g. cell death, molecular markers, or behavioural changes). | Yes |
| | | b. For hypothesis-testing studies, specify the primary outcome measure, i.e. the outcome measure that was used to determine the sample size. | Yes |
| **Statistical methods** | 7 | a. Provide details of the statistical methods used for each analysis, including software used. | Yes |
| | | b. Describe any methods used to assess whether the data met the assumptions of the statistical approach, and what was done if the assumptions were not met. | Yes |
| **Experimental animals** | 8 | a. Provide species-appropriate details of the animals used, including species, strain and substrain, sex, age or developmental stage, and, if relevant, weight. | Yes |
| | | b. Provide further relevant information on the provenance of animals, health/immune status, genetic modification status, genotype, and any previous procedures. | Yes |
| **Experimental procedures** | 9 | For each experimental group, including controls, describe the procedures in enough detail to allow others to replicate them, including: | Yes |
| | | a. What was done, how it was done and what was used. | Yes |
| | | b. When and how often. | Yes |
| | | c. Where (including detail of any acclimatisation periods). | |
| | | d. Why (provide rationale for procedures). | Yes |
| **Results** | 10 | For each experiment conducted, including independent replications, report: | |
| | | a. Summary/descriptive statistics for each experimental group, with a measure of variability where applicable (e.g. mean and SD, or median and range). | Yes |
| | | b. If applicable, the effect size with a confidence interval. | Yes |
